# Inference and visualization of phenome-wide causal relationships using genetic data: an application to dental caries and periodontitis

**DOI:** 10.1101/865956

**Authors:** Simon Haworth, Pik Fang Kho, Pernilla Lif Holgerson, Liang-Dar Hwang, Nicholas J. Timpson, Miguel E. Rentería, Ingegerd Johansson, Gabriel Cuellar-Partida

## Abstract

**Background:** Hypothesis-free Mendelian randomization studies provide a way to assess the causal relevance of a trait across the human phenome but can be limited by statistical power or complicated by horizontal pleiotropy. The recently described latent causal variable (LCV) approach provides an alternative method for causal inference which might be useful in hypothesis-free experiments.

**Methods:** We developed an automated pipeline for phenome-wide tests using the LCV approach including steps to estimate partial genetic causality, filter to a meaningful set of estimates, apply correction for multiple testing and then present the findings in a graphical summary termed a causal architecture plot. We apply this process to body mass index and lipid traits as exemplars of traits where there is strong prior expectation for causal effects and dental caries and periodontitis as exemplars of traits where there is a need for causal inference.

**Results:** The results for lipids and BMI suggest that these traits are best viewed as creating consequences on a multitude of traits and conditions, thus providing additional evidence that supports viewing these traits as targets for interventions to improve health. On the other hand, caries and periodontitis are best viewed as a downstream consequence of other traits and diseases rather than a cause of ill health.

**Conclusions:** The automated process is available as part of the MASSIVE pipeline from the Complex-Traits Genetics Virtual Lab (https://vl.genoma.io) and results are available in (https://view.genoma.io). We propose causal architecture plots based on phenome-wide partial genetic causality estimates as a way visualizing the overall causal map of the human phenome.

**Key messages:**

1. The latent causal variable approach uses summary statistics from genome-wide association studies to estimate a parameter termed *genetic causality proportion*.
2. Systematic estimation of genetic causality proportion for many pairs of traits provides an alternative method for phenome-wide causal inference with some theoretical and practical advantages compared to phenome-wide Mendelian randomization.
3. Using this approach, we confirm that lipid traits are an upstream risk factor for other traits and diseases, and we identify that dental diseases are predominantly a downstream consequence of other traits rather than a cause of poor systemic health.
4. The method assumes no bidirectional causality and no confounding by environmental correlates of genotypes, so care is needed when these assumptions are not met.
5. We developed an automated and accessible pipeline for estimating phenome-wide causal relationships and generating interactive visual summaries.

## Introduction

Associations between causal risk factors and disease can suggest new ways to improve health. Conventional epidemiological studies may uncover correlations but cannot easily disentangle non-causal or reverse-causal relationships where interventions on the putative risk factor will be ineffective. In this article, risk factors are described as “upstream” if they have effects on disease, or “downstream” if the putative risk factor is a marker of or a consequence of the disease, irrespective of chronology.

Dental diseases are good examples of complex diseases which are associated with a range of poor health outcomes and are hypothesized to be both a cause and consequence of ill health^1^ but the limitations of conventional epidemiological methods do not exclude confounded association. In the context of recent calls to prioritize prevention and early interventions, address the global health problem of dental diseases and overcome isolation between dentistry and medicine^2, 3^, there is a need to locate dental diseases in the context of causal flow through the human phenome. Conversely, lipid biomarkers such as low-density lipoprotein cholesterol (LDL-C) are good examples of traits which are known to have effects on human health including cardiovascular disease^4, 5^ and may act as a positive control for contemporary epidemiological methods which aim to identify causal relationships.

In recent years techniques have been proposed which use genetic data to try and assert causality in observational studies^6^ and these are particularly valuable in situations where large scale interventional studies would be impractical or unethical. One example is Mendelian Randomization (MR), an analytical paradigm which uses genetic variants as proxies for a risk factor in order to test for causal effects on an outcome^7^. In dental epidemiology, this method has been used to examine the effects of potentially modifiable risk factors like Vitamin D and body mass index on caries and periodontitis^8, 9^, to assess the possible impact of periodontitis on hypertension^10^ and undertake bi-directional analysis to test for causal relationships between dental diseases and cardio-metabolic traits in both directions^11^. To date, these studies have only explored a small number of traits and the bespoke experimental design used for each study makes it difficult to compare estimates for different diseases. Dental diseases may therefore serve as a model for complex traits where it would be helpful to perform a causal inference analysis in a systematic manner across the whole phenome.

There are practical challenges meaning that MR may not be the preferred approach for a phenome-wide causal experiment in this context. At its heart, MR experiments rely of vertical pleiotropy, that is to say a genotype with effects on trait A is associated with trait B because trait A affects trait B. It can be difficult to distinguish this from horizontal pleiotropy, where a genetic variant has biological effects on both trait A and trait B. Many genetic variants have horizontally pleiotropic effects, leading to false positive findings or over-estimation in effect sizes at true positive associations in classical MR experiments^11, 12^. Several estimation techniques have been developed which use the distribution of causal effect estimates across multiple variants in an attempt to detect and account for^13-15^ or at least reduce the impact of horizontal pleiotropy^16^. These methods may however introduce additional assumptions about the distribution of effect estimates^17, 18^ and run into problems when these assumptions are not met^19^, suggesting each estimate produced using these methods may need interpretation on a case by case basis to assess whether the assumptions are reasonable. In addition, MR experiments can produce spurious findings due to sample overlap ^20^ which can be problematic in phenome-wide studies, as the same underlying population in a consortium or biobank may contribute to the available genetic evidence for many different traits. Finally, MR experiments use information from a small number of genetic variants and discard information from most of the genome, meaning that statistical power may be limited for a phenome-wide experiment for traits such as dental diseases which have relatively few robust single variant association signals.

An alternative analytical paradigm – the latent causal variable (LCV) method – has recently been proposed. LCV uses information aggregated across the whole genome to infer potentially causal relationships between complex human traits and diseases^21^. In conjunction with large-scale genetic association studies made possible by resources such as UK Biobank^22^ and automated pipelines for quality control and analysis such as the Complex-Traits Genetics Virtual Lab (CTG-VL)^23^, this method now provides an opportunity to obtain information on potentially causal relationships efficiently and at phenome-wide scale. Here we introduce CTG-VL’s newly implemented capability to perform a phenome-wide scan across hundreds of traits using the LCV method and the visualization of the results using causal architecture plots. We showcase this method using GWAS data for body mass index, lipid levels, dental caries and periodontitis^11^.

## Methods

### Conceptual overview

The genetic correlation between two traits represents the correlation in genetic effect sizes at common genetic variants across traits^24^. The latent causal variable (LCV) approach initially estimates the genetic correlation between traits A and B using a modified linkage disequilibrium score regression technique, which can detect and account for sample overlap in genetic association studies^24^, and subsequent stages are only informative when there is detectable genetic correlation. Next, the model fits a single unobserved variable (termed *L*) which is causal for trait A and trait B and mediates the observed genetic correlation between traits A and B. To distinguish between horizontal and vertical pleiotropy the LCV model compares the correlation between *L* and trait A with the correlation between *L* and trait B and estimates a parameter termed genetic causality proportion (GCP). Positive values of GCP suggest vertical pleiotropy where trait A lies upstream of trait B and interventions on trait A are likely to affect trait B while negative GCP values indicate that B lies upstream of trait A. GCP values near 0 imply that the genetic correlation between traits A and B is mediated by horizontal pleiotropy and interventions on traits A or B are unlikely to affect the other trait. A detailed description of the LCV method is provided in the original publication^21^.

### Using the distribution of GCP estimates to infer the causal architecture of a trait

If traits A and trait B are swapped the GCP estimate is unchanged in magnitude however the sign is reversed. In an experiment involving all pairwise comparisons between *n* traits this creates symmetry, which is to say for every positive signed GCP estimate observed in the experiment there must be an equal but negatively signed GCP estimate corresponding to the same pair of traits but with the order of traits reversed. If a randomly selected trait from group *n* has predominantly positive GCP estimates, this implies that the trait is an upstream factor of the majority of other traits in group *n.* Conversely, if the GCP estimates are predominantly negative, this implies that the trait is a downstream factor of most other traits in group *n* and interventions on this trait are less likely to change the other traits in group *n.*

We suggest that if two or more target traits are compared against the same panel of anchor traits, then differences in the distribution of GCP estimates between those traits provide an indication of which target traits may have a greater or lesser causal relevance (assuming GWAS of anchor traits are equally powered – see Discussion) for the human phenome, which traits represent upstream determinants of health and which are downstream consequences of other traits. We propose an automated pipeline for obtaining GCP estimates for target traits against a shared panel of anchor traits and visualizing the results in a causal architecture plot.

### Pipeline stages and implementation

All traits conducted in studies of European ancestry in CTG-VL catalogue were selected. CTG-VL is a curated resource of genome-wide association (GWA) summary statistics and downstream analysis ^23^. The complete list of GWA summary statistics and references are available in CTG-VL. Briefly, these data were derived from various international genetics consortia and UK Biobank, where the inclusion criteria was a nominally significant (P<0.05) single nucleotide polymorphism (SNP) based heritability derived from LD-score regression^25^. In total 1389 studies are currently available in CTG-VL. References for each of these GWAS are directly available in CTG-VL.

#### Traits selection

As a positive control, the analyses were first performed on GWAS summary statistics for high density lipoprotein cholesterol (HDL-C, n=188,577), low density lipoprotein-cholesterol (LDL-C, n=188,578), total cholesterol (TC, n=188,579), triglycerides (TG, n=188,580) ^26^ and body mass index (BMI) (n=339,224) ^27^ where we expected to observe effects of these traits on a multitude of traits and conditions. We then showcase this pipeline using GWAS summary statistics of dental caries and periodontitis due to a paucity of existing causal evidence. Genetic association data for dental disease traits were taken from genome-wide association studies which combined clinical data from the GLIDE consortium with genetically validated proxy phenotypes from UK Biobank as previously described^11^. Data were combined using a z-score genome-wide meta-analysis weighted by effective sample size. The traits were a) decayed, missing and filled tooth surfaces (n=26,792 from nine studies in GLIDE) and dentures (n_cases_=77,714, n_controls_=383,317 in UK Biobank) and b) periodontitis (n_cases_=17,353, n_controls_=28,210 from seven studies) and loose teeth (n_cases_=18,979, n_controls_=442,052).

#### Analysis

The R version (URL: https://github.com/lukejoconnor/LCV) for the LCV method made available by the original authors ^21^ was implemented in CTG-VL to carry out phenome-wide scans as parts of CTG-VL’s MASSIVE (*Massive downstream analysis of summary statistics*) pipeline (URL: https://vl.genoma.io). LCV models were fitted in a pairwise manner comparing each test trait to each anchor trait using the automated implementation of the LCV method in CTG-VL. For the analysis we used the subset of genetic variants present in the HapMap3 consortium dataset ^28^ and linkage disequilibrium scores obtained from European ancestry samples within the 1000 genomes project data (phase 3, 2018 release, provided by provided by Alkes Price’s group (URL: https://data.broadinstitute.org/alkesgroup/LDSCORE/).

#### Post-processing

LCV estimates are only informative when there is evidence for genetic correlation between the target trait and anchor trait. First, traits with evidence for a non-zero genetic correlation (Benjamini-Hochberg’s FDR < 5%) were carried forward. Next, we ran LCV analyses to estimate GCP in the remaining traits and again applied a Benjamini-Hochberg’s FDR < 5% to the GCP p-value (H_0_: GCP =0).

#### Causal architecture plots

To visualize a target trait in the context of the human phenome we propose a visual summary termed a causal architecture plot (Figure 2). Each marker indicates an anchor trait where there is detectable genetic correlation with the target trait so plots with a complex target trait with few markers may indicate low heritability or an underpowered GWAS. The Y axis represents the strength of evidence for causal relationship between the target trait and anchor trait with a red line indicating which relationships pass multiple test correction, allowing differences between traits with limited causal relevance or widespread causal relevance to be identified. A symmetrical funnel plot indicates equal numbers of upstream and downstream factors for the target trait (e.g. Figure 2A, Figure 2D), while an asymmetrical funnel indicates the causal direction is predominantly from the anchor traits to the target trait (e.g. Figure 2E) or from the target trait to the anchor traits (Figure 2E), as shown by the different shaded zones (Figure 2F). The markers are colored to show the strength and direction of genetic correlation, which also indicates whether causal relationships are in a trait-increasing or trait-decreasing direction. Finally, the size of markers provides an indication about the precision of the LCV estimates and labels provide the names of the anchor traits with strongest evidence for causal association.

**Figure 1:**
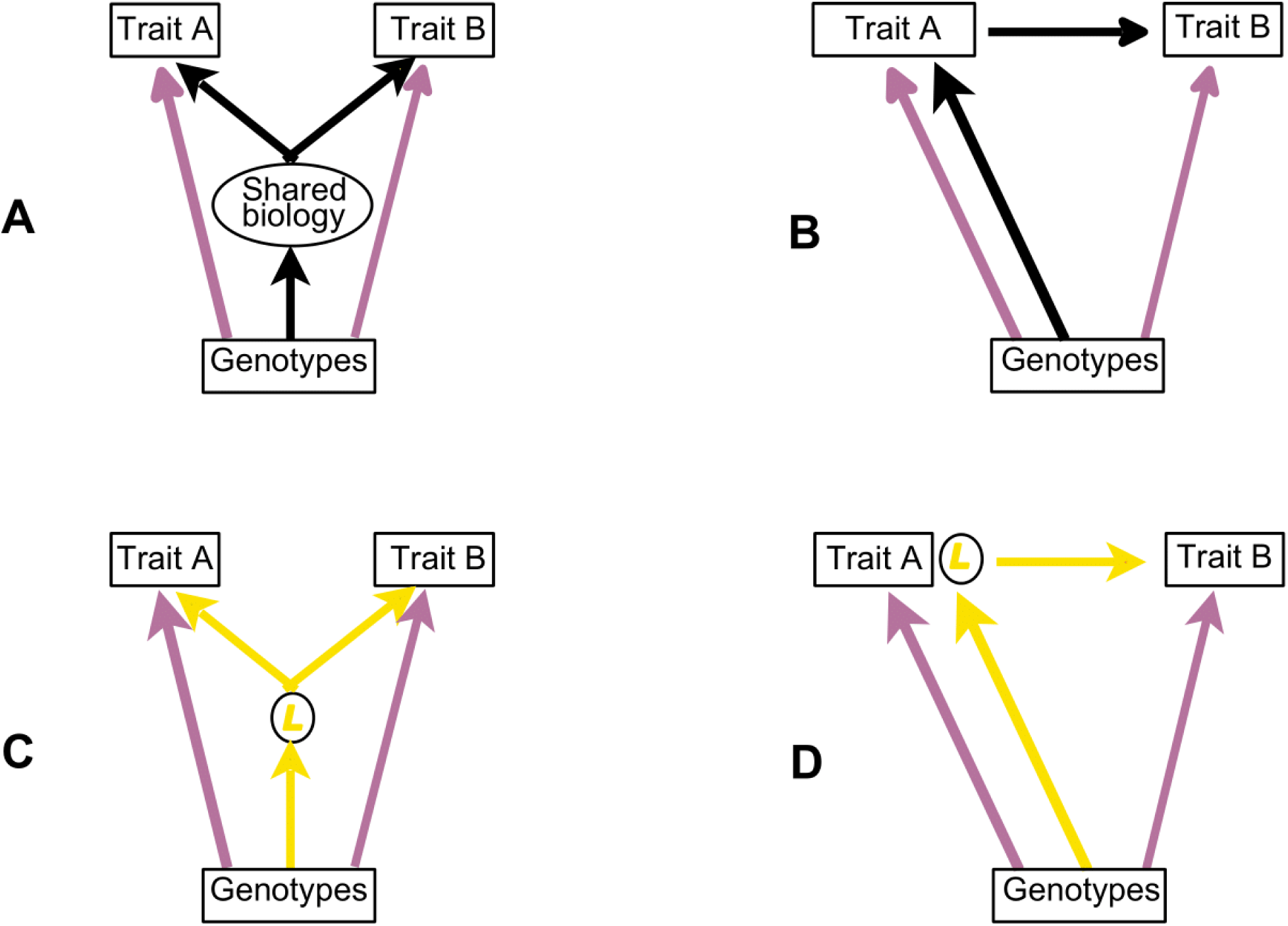
Use of a latent variable for causal inference. The genetic effects on trait A and B (purple arrows) are correlated in all scenarios. This could be due to horizontal pleiotropy (1A, true causal pathway drawn in black), vertical pleiotropy (1B, true causal pathway drawn in black) or a combination of both processes. In LCV analysis an inferred causal pathway is created which mediates the observed genetic correlation between traits A and B and must always pass through *L.* Where horizontal pleiotropy mediates the genetic correlation between traits A and B, the genetic correlation between *L* and traits A and B is similar in magnitude giving a GCP estimate near zero (Figure 1C, inferred causal pathway drawn in yellow, true causal pathway shown in 1A). In situations where *L* has a perfect genetic correlation with trait A, the only effects of genotypes on trait B must be through their effects on trait A, (1D, inferred causal pathway drawn in yellow, true causal pathway shown in 1B), analogous to a positive finding in a classical MR experiment using a valid instrument and resulting in a GCP estimate of 1.

**Figure 2:**
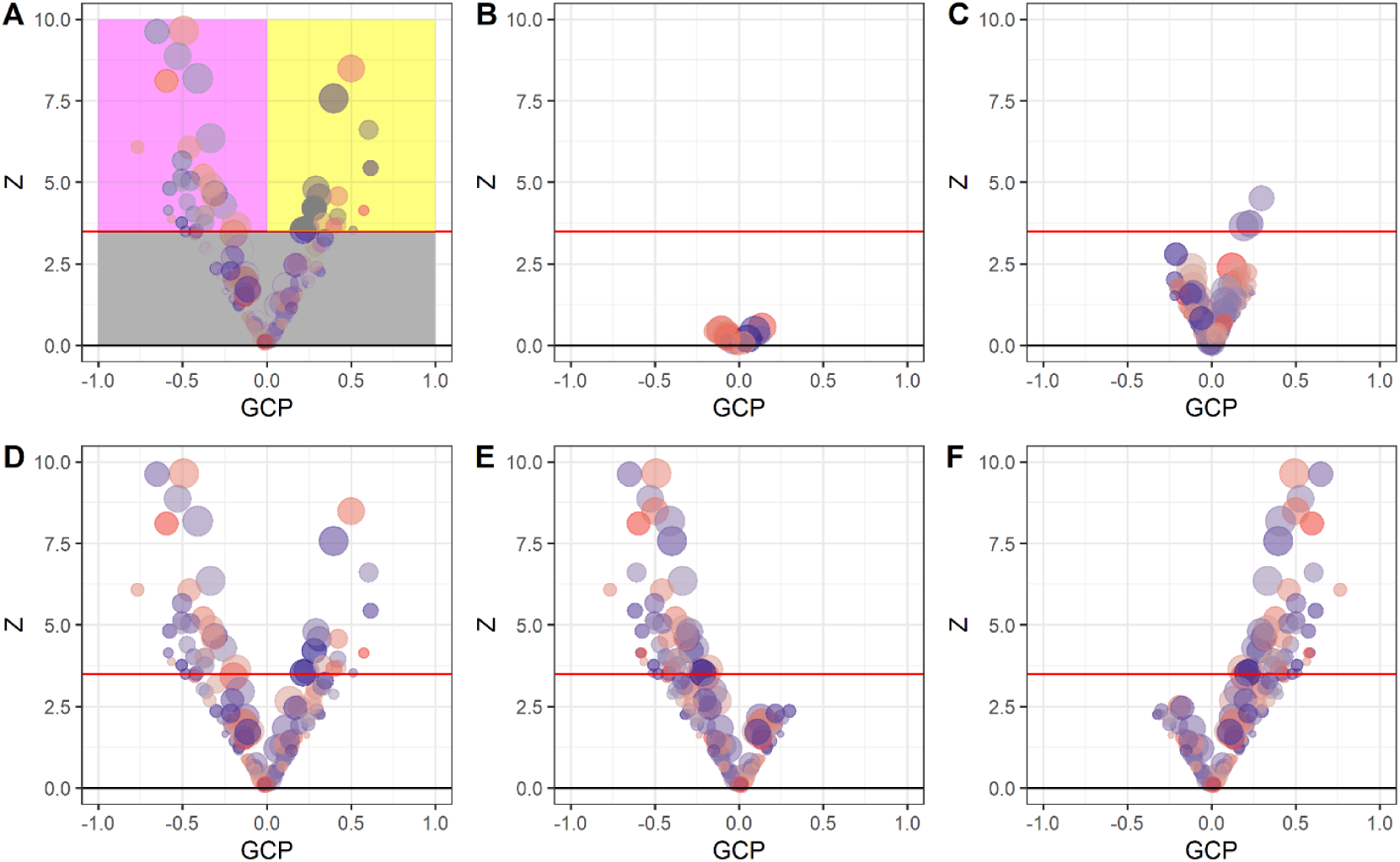
Interpretation of causal architecture plots. Each dot represents a test trait tested against the anchor trait and the red line represents the statistical significance threshold (FDR < 5%). A) Schematic showing regions of the plot which represent non-causal relationships (grey), upstream (pink) and downstream (yellow) causal relationships, B) Under-powered experiment, C) well-powered experiment for a trait with limited causal relevance, D) The trait has many causal relationships in both upstream and downstream directions, E) A trait which is causally affected by other upstream traits F) A trait with downstream effects.

## Results

### Lipid traits

We observed that LDL, HDL, TG and TC produced causal architecture plots showing only downstream effects on several traits (Figure 3). Table 1 summarizes the number of causal relationships estimated by LCV per each trait and Supplementary Table 1-4 show the complete list of results. HDL had many causal relationships in a trait-decreasing (risk reducing for disease traits) direction while effects of TG were predominantly in a trait-increasing direction. TC and LDL had relatively few genetic correlations however a large proportion of these were partially due to causal effects, again, predominantly in a downstream direction.

**Table 1:** Summary of the results of the LCV method for each trait.

| Trait | Number of genetic correlations with FDR < 5%. | Number of potentially causal relationships (FDR < 5%) | Upstream/downstream traits |
| --- | --- | --- | --- |
| HDL | 253 | 140 | 2/140 |
| LDL | 14 | 12 | 0/12 |
| TC | 23 | 18 | 0/18 |
| TG | 186 | 96 | 2/94 |
| BMI | 647 | 133 | 23/110 |
| Caries | 527 | 71 | 71/0 |
| Periodontitis | 398 | 32 | 29/3 |

\*Each trait was tested against the same panel of 1,389 partially heritable traits in CTG-VL catalogue. All the results with FDR < 5% are provided in Supplementary Table 1-7 and full results are browsable using CTG-View platform (URL: https://view.genoma.io).

**Figure 3:**
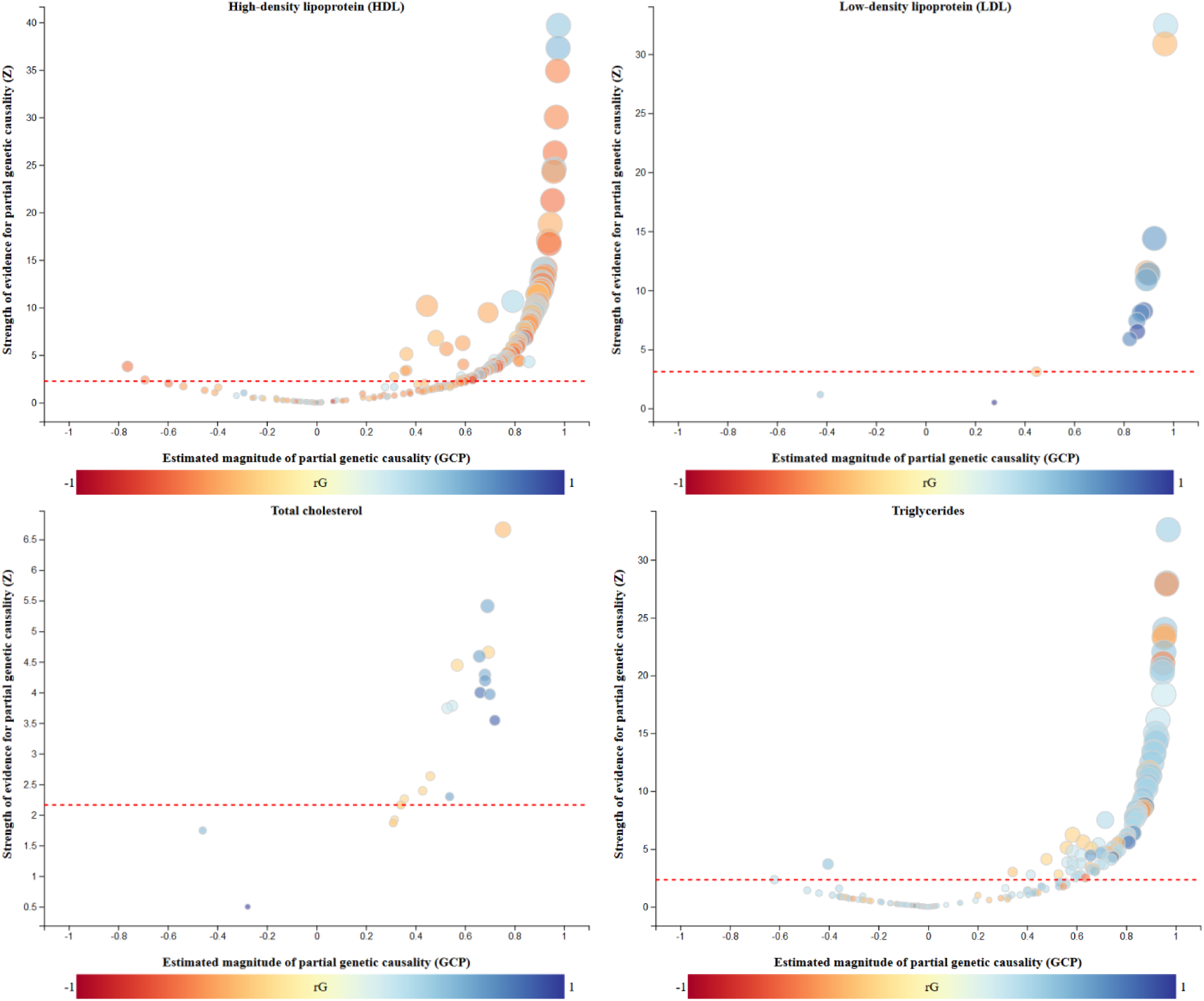
Comparison of the causal architecture of lipid traits. Only traits with a genetic correlation at FDR < 5% are shown. The red dashed line corresponds to the statistical significance threshold (FDR < 5%) for genetic causality proportions.

### BMI, caries and periodontitis

In part, the ability of LCV to resolve clear differences between the four lipid traits might be helped by the relatively simple genetic architecture of these traits. By contrast, complex traits such as BMI which are affected by many different biological processes may provide a more realistic control for comparison against caries and periodontitis.

For BMI, genetic correlations with 647 traits were identified, of which 133 were partially due to causal relationships. The majority of GCP estimates were positively signed, suggesting that BMI may impact many other traits (Figure 4A) however there were also several negatively signed relationships, suggesting that BMI it itself could potentially be amenable to several different interventions. The upstream trait with greatest evidence for causal effect was a BMI-increasing effect of employment as a heavy goods vehicle driver, while the downstream trait with greatest evidence was ‘vascular/heart problems diagnosed by doctor’, where a lower BMI was associated with greater odds of reporting no vascular or heart problems (Figure 4A and Supplementary Table 5).

**Figure 4.**
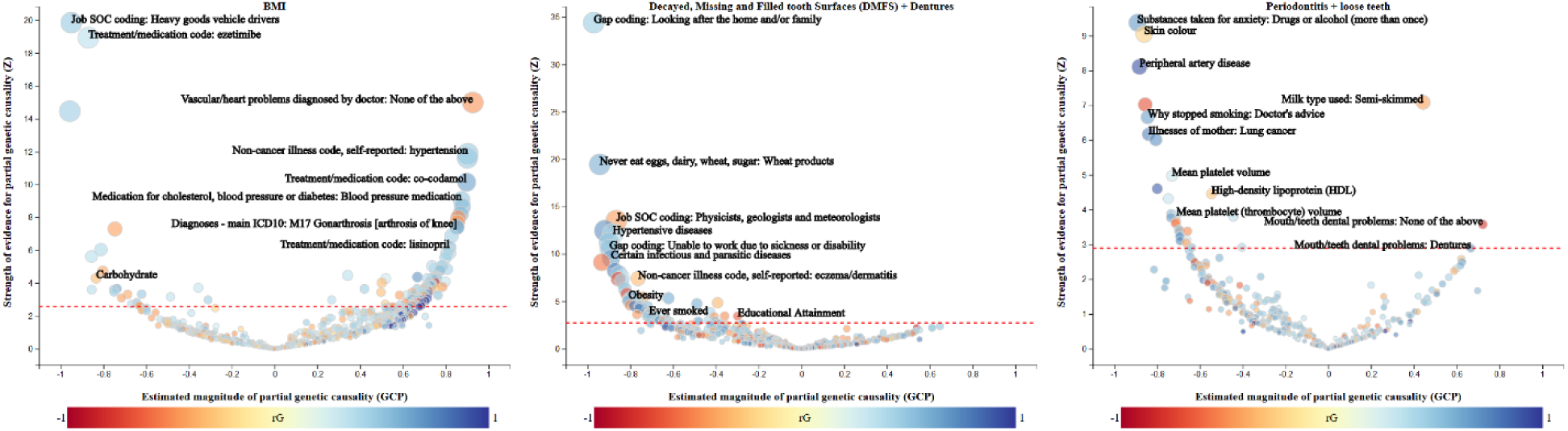
Comparison of the causal architecture of BMI, caries and periodontitis. BMI, B) Caries proxied by DMFS/dentures, C) Periodontitis proxied by periodontitis/loose teeth. All the statistically significance results (FDR < 5%) are shown in Supplementary Tables 5-7 and can be queried at https://view.genoma.io.

For dental caries proxied by DMFS/dentures there were detectable genetic correlations with 527 traits, of which 71 supported a partially causal relationship (Figure 4B). All GCP estimates were negatively signed, suggesting that DMFS/dentures is a downstream consequence of these traits rather than a causal risk factor. Traits with evidence for partial genetic causality included harmful effects of variables capturing dietary habits, smoking, hypertensive diseases and obesity while a protective effect was observed for variable representing skilled employment and education (Figure 4B and Supplementary Table 6).

For periodontitis proxied by the combination of periodontitis/loose teeth 398 genetic correlations were detected at FDR < 5%, of which a relatively small faction (32) were modelled to be partially due to causal relationships. The predominant direction was negatively signed (29 out of 32 traits), again suggesting the predominant direction of causality is from other traits to periodontitis (Figure 4C). The 5 traits with the strongest evidence for partial genetic causality were a) a harmful effect of drug or alcohol use for anxiety on periodontitis b) a protective effect of fairer skin color on periodontitis c) a harmful effect of peripheral artery disease on periodontitis, d) an effect of periodontitis on dietary preference (proxied by preferred type of milk) and e) a protective effect of a variable representing absence of problematic alcohol consumption. Periodontitis appeared to have a causal effect with other dental problems and increase in the use of dentures.

## Discussion

Previous approaches to obtaining phenome-wide causal maps have been based around the Mendelian Randomization paradigm^29^. The LCV method has attractive properties for phenome-wide analysis as it is robust to sample overlap, has greater statistical power than MR^17^ and is unconfounded by horizontal pleiotropy^21^. We developed a pipeline to automate LCV analysis and visualize results in causal architecture plots and applied this to lipid traits and BMI as positive controls, and to caries and periodontitis as exemplars of complex traits where there is a need for additional causal evidence. The results suggest that, at a high level, dental diseases are embedded in the human phenome but best viewed as a downstream marker of biological events and a consequence of other diseases rather than as a driver of biological changes which lead to large or widespread changes in other traits. The results therefore support the current drive to target upstream determinants of dental diseases^3^ and potentially provide a framework for prioritizing subsets of traits which have greater or reduced causal relevance for detailed epidemiological analysis or translational research. Specifically, the results for caries and periodontitis prioritize socio-economic status, cardiovascular health, diet and mental health/alcohol use as traits which could be targeted to improve dental health. Conversely, the results for HDL-C confirm that interventions on HDL-C are likely to have downstream effects on many traits and diseases, and that BMI is a trait with many causal relationships in both upstream and downstream directions. We suggest that this pipeline may be helpful to researchers undertaking initial characterization of a phenotype, and have implemented it as part of CTG-VL, a freely available online resource (URL: https://vl.genoma.io)

The LCV method requires GWA summary statistic data and needs to identify a genetic correlation between the target trait and anchor trait for the results to be meaningful. It was therefore only possible to examine traits which have been studied using a large enough GWAS to yield a stable heritability estimate. While this captures many important diseases, risk factors and intermediate traits reflected by the large number of anchor traits, there are natural limitations to the results which are available at this moment in time. For example, risk factors or outcomes such as the oral microbiota composition, oral health quality of life, dental anxiety and satisfaction with dental appearance and function may be causally related to dental diseases but are not represented by current genome-wide association studies. For dental diseases specifically, this illustrates the need to ensure that oral and dental health is represented in epidemiological studies using current methods to avoid perpetrating the under-representation of dentistry in the next generation of epidemiological research. As the number of curated GWAS summary statistics in CTG-VL catalogue increases over time, this limitation will become less important. It will become possible to construct more detailed causal architecture plots for any given target trait, and it may be possible to move from single-trait causal profiles towards multi-trait visualization which present an overall causal map of the human phenome.

While this pipeline is primarily intended to give an overview of a trait, it may also highlight specific findings which warrant further investigation. One interesting pair of findings were that hair colour appears to be an upstream determinant of dental caries and that skin colour appears to be a risk factor for periodontitis. These findings may have a biological explanation (for example both ancestry and skin colour are associated with periodontitis in observational studies^30, 31^, skin color is associated with caries in children with a possible mechanism related to vitamin D^32^ and hair keratins have a role in enamel formation which might predispose to caries^33, 34^). Alternatively, the findings may also reflect complexity introduced by the scale and sampling frame of UK Biobank. Although the LCV model is more robust than MR to biasing effects from sources such as horizontal pleiotropy and sample overlap, the LCV model can become biased by correlation between genetic variation and environmental factors which affect disease^17, 35^. This aggregation might be due to factors such as ancient ancestry^36^, genetic nurture effects^37^ or sampling phenomena^38^ and is a concern in the UK Biobank sample^39^ where much of the data used in this experiment were obtained. Interpreted in this light, it is possible that environmental factors which are more prevalent in groups of people with certain hair type or skin color are a cause of dental diseases. This example may therefore illustrate some of the challenges created by population stratification but also the opportunities for genetic information to inform research about social and environmental factors which may affect disease.

Previous studies using the MR method have found some evidence for causal effects of caries and periodontitis on cardiovascular health traits^10, 11^ which was not recapitulated using the LCV method. In part, this may be because LCV aims to captures the overall or predominant direction of causality mediated by a single latent variable and may therefore be a poor fit to systems with complex features such as polytonicity, non-linear effects or bidirectional causality. We suggest that the causal architecture plots are used to provide an overall causal context to a trait as an adjunct to other methods to assert causality which have different strengths and limitations. Despite this, profiles for lipid traits were obtained under the same analytical conditions but appear strikingly different, providing a clear indication that the method can resolve major differences in causal architecture between diseases.

It is important to recognize some limitation of this work. Here, we presented a pipeline to do a phenome-wide scan of potential causal associations. However, it is important to note that the current set of GWAS do not encompass the complete phenome and this is biased towards well powered GWAS and thus restricted to common diseases and traits. As the range of GWAS studies increases with time, this limitation will become less important. It is also important to recognize that GCP estimates are also tied to the statistical power of the GWAS, thus impacting the ability to detect causal effects for specific traits. Low statistical power of GWAS does not however bias the model towards positive or negative values of GCP, so the split of upstream/downstream estimates will still be informative even when there are relatively few causal effects identified. Finally, the model assumes that the GWAS for both traits and reference LD data are drawn from the same underlying population, which at present limits this pipeline to analysis of studies of European ancestry participants. As the number of GWAS studies in diverse populations increases and additional reference datasets become available, it may be possible to extend this method to other populations.

In summary, we present a pipeline to estimate and visualize genetic causality proportions across traits with GWAS summary statistics implemented in CTG-VL. All the results are freely available for download in https://view.genoma.io.

## Acknowledgements

SH is funded by a National Institute for Health Research (NIHR) Academic Clinical Fellowship. GCP is funded by an Australia Research Council Discovery Early Career Researcher Award (DE180100976).

## Data access

No participant-level data were accessed to produce this article. The sources of GWA summary statistics and reference used to perform analysis are described in full in the methods.

